# Atypical functional connectivity in adolescents and adults with persistent and remitted ADHD

**DOI:** 10.1101/201772

**Authors:** Giorgia Michelini, Joseph Jurgiel, Ioannis Bakolis, Celeste H. M. Cheung, Philip Asherson, Sandra K. Loo, Jonna Kuntsi, Iman Mohammad-Rezazadeh

## Abstract

We previously provided initial evidence for cognitive and event-related potential markers of persistence/remission of attention-deficit/hyperactivity disorder (ADHD) from childhood to adolescence and adulthood. In this follow-up study, using a novel brain-network connectivity approach, we aimed to examine whether functional connectivity reflects a marker of ADHD remission, or an enduring deficit unrelated to ADHD outcome. High-density EEG was recorded in 110 adolescents and young adults with childhood ADHD (87 persisters, 23 remitters) and 169 typically-developing individuals during an arrow-flanker task, eliciting cognitive control. Functional connectivity was quantified with network-based graph-theory metrics before target onset (pre-stimulus), during target processing (post-stimulus) and in the degree of change between pre-stimulus/post-stimulus. ADHD outcome was examined with parent-reported symptoms and impairment using both a categorical (DSM-IV) and a dimensional approach. Graph-theory measures converged in indicating that, compared to controls, ADHD persisters showed increased connectivity in pre-stimulus theta, alpha and beta and in post-stimulus beta (all p<.01), and reduced pre-stimulus/post-stimulus change in theta connectivity (p<.01). In the majority of indices showing ADHD persister-control differences, ADHD remitters differed from controls (all p<.05), but not from persisters. Similarly, connectivity measures were not associated with continuous outcome measures of ADHD symptoms and impairment in participants with childhood ADHD. These findings indicate that adolescents and young adults with persistent and remitted ADHD share atypical over-connectivity profiles and reduced ability to modulate connectivity patterns with task demands, compared to controls. Brain connectivity impairments may represent enduring deficits in individuals with childhood ADHD irrespective of diagnostic status in adolescence/young adulthood.

## INTRODUCTION

A coherent communication between different brain regions, or brain functional connectivity, is thought to have a key role in cognition and behavior^1-3^. Accumulating evidence suggests that atypical connectivity may be implicated in neurodevelopmental disorders^4-6^, such as attention-deficit/hyperactivity disorder (ADHD). Most studies to date have investigated brain connectivity in ADHD using functional magnetic-resonance imaging (fMRI), with reduced connectivity within and between brain regions/sub-networks during resting (e.g., within the default-mode network (DMN) and between DMN and executive networks) observed in individuals with ADHD^7-11^. Evidence of increased resting-state connectivity within and between these regions, however, has also been reported in ADHD^4, 12-16^. Examining brain connectivity during task performance further allows a more direct characterization of connectivity alterations underlying the impairments in cognition and behavior associated with ADHD^17, 18^. Task-based fMRI studies of ADHD show hypo-connectivity in fronto-striato-cerebellar networks during sustained attention^19^ and inhibition^20-22^, and hyper-connectivity within the DMN^20^ and between networks of reward/cognitive control integration^23^. Using the sub-second temporal resolution of electroencephalography (EEG), previous studies have further shown hypo- and hyper-connectivity in slower and faster brain oscillations from different cortical regions during rest in individuals with ADHD^24-26^. Available task-based studies in children and adolescents with ADHD indicate reduced fronto-parietal theta-alpha connectivity^27, 28^, but also increased connectivity in alpha^29^ and beta^30^. No study to date has examined task-based EEG connectivity in adults with ADHD. Overall, despite inconsistencies regarding which brain networks may be hypo- and hyper-connected, available evidence points to atypical brain connectivity in ADHD.

While atypical functional connectivity has been documented both in children^7-9^ and adults^10, 11, 31^ with ADHD, little is known on how these alterations map onto ADHD developmental outcomes. ADHD persists, in full or in partial remission, in the majority of adolescents and adults clinically diagnosed in childhood^32, 33^. Yet, the evidence that some individuals remit across development may suggest the presence of (1) neural processes that are markers of remission, improving concurrently with clinical profiles and distinguishing individuals with persistent and remitted ADHD (ADHD “persisters” and “remitters”, respectively); and of (2) enduring deficits that are unrelated to the clinical outcome, remaining impaired in both remitters and persisters^34^. The identification of such measures may help elucidate the mechanisms underlying remission/persistence, and point to candidate biomarkers for the development of new interventions for ADHD. Most studies to date, using cognitive-performance indices, found that executive functioning measures do not distinguish between ADHD persisters and remitters, and are thus insensitive to ADHD outcomes^35-39^. Fewer studies have investigated the neural underpinnings of remission/persistence. In a recent follow-up of adolescents and young adults with childhood ADHD, we found that cognitive and event-related potential (ERP) markers of executive control (inhibition, working memory, conflict monitoring) were insensitive to ADHD outcome^38-40^. Instead, cognitive measures and EEG activity of preparation-vigilance and error detection were markers of remission, distinguishing ADHD remitters from persisters.

Considering the important role of brain connectivity in behavior and cognition^1-3^, investigating this brain-wide neural mechanism may provide new insight into the neural pathways of ADHD persistence/remission. Only three studies to date have examined functional connectivity in remitted and persistent ADHD, using fMRI^31, 41, 42^. Two of these studies, using small samples, reported that ADHD persisters showed lower functional connectivity than remitters and controls between the DMN and executive network during rest^31^ and between the thalamus and frontal areas during response preparation^41^. Resting-state medial-dorsolateral functional associations in the prefrontal cortex, implicated in cognitive control, was instead unrelated to ADHD outcome, and reduced in both ADHD remitters and persisters, compared to controls^31^. Another study, however, found higher connectivity in ADHD remitters than controls, with persisters showing intermediate profiles between remitters and controls^42^. Investigating brain connectivity using the excellent temporal resolution of EEG may provide further information in relation to ADHD remission/persistence by capturing fast and transient changes in functional connectivity (not captured by fMRI) during cognitive processes^43, 44^. Yet, most EEG connectivity studies in ADHD to date present methodological limitations, such as the use of connectivity metrics contaminated by volume-conduction artefacts (i.e., the spreading and mixing of multiple brain sources at the scalp), which may produce inflated connectivity estimates^45, 46^. Recently developed network approaches, such as graph theory, may be further applied to characterize brain connectivity between large-scale brain networks and identify connectivity alteration^2, 4, 47^. Initial graph-theoretic evidence from two task-based studies shows atypical functional connectivity in children with ADHD^48, 49^. No study to date has examined EEG connectivity in relation to longitudinal ADHD outcome.

In the present EEG study, we aimed to investigate brain functional connectivity during a cognitive control task, the arrow flanker task, in a follow-up of adolescents and adults with childhood ADHD and neurotypical controls. In previous analyses on the present sample with this task, we have shown that attention-vigilance cognitive processes (reaction-time variability [RTV] and errors in low-conflict trials) and ERPs of error detection (error-related negativity [ERN] and positivity [Pe] amplitudes) were markers of remission, as ADHD remitters differed from persisters but not from controls in these measures^39^. Instead, cognitive-ERP measures of executive and conflict processes (errors in high-conflict trials, N2 amplitude) and processing speed (mean reaction time [MRT]), did not distinguish ADHD remitters from persisters, despite being sensitive to differences between persisters and controls^39^. Here, we aimed to test whether functional connectivity, measured with graph-theory and connectivity metrics not contaminated by volume conduction, is atypical in persistent ADHD, and whether it represent a marker of ADHD remission or an enduring deficit. We hypothesized that both ADHD persisters and remitters would display functional connectivity alterations compared to neurotypical individuals during this task evoking high levels of cognitive control, consistent with most studies examining cognitive and EEG markers of executive processes and ADHD remission^35-39^.

## METHODS

### Sample

The sample consisted of 279 participants who were followed up on average 5.8 years (SD=1.1) after assessments in childhood^50^, including 110 adolescents and young adults who met DSM-IV criteria for combined-type ADHD in childhood (10 sibling pairs and 90 singletons) and 169 control participants (76 sibling pairs and 17 singletons)^38, 51^. Participants with ADHD were initially recruited from specialized ADHD clinics, and controls from schools in the UK^50^. Exclusion criteria at both assessments were: IQ<70, autism, epilepsy, brain disorders, and any genetic/medical disorder associated with externalizing behaviors that might mimic ADHD. Among those with childhood ADHD, at follow-up 87 (79%) continued to meet clinical (DSM-IV) levels of ADHD symptoms and impairment (ADHD persisters), while 23 (21%) were below the clinical cut-off (ADHD remitters)^52^. Among ADHD remitters, 14 displayed ≥5 symptoms of inattention or hyperactivity-impulsivity, but did not show functional impairment. Participants attended a single research session for clinical, IQ and cognitive-EEG assessments. An estimate of IQ was derived with the vocabulary and block design subtests of the Wechsler Abbreviated Scale of Intelligence (WASI)^53^. ADHD persisters, remitters and controls did not differ in age, but there were significantly more males in the remitted group than in the other two groups, with no females among ADHD remitters (Table 1)^38, 39^. ADHD persisters showed lower IQ compared to remitters and controls^38, 52^. 47% of participants with childhood ADHD were on drug treatment at follow-up, but the proportion of participants on medication did not differ between ADHD persisters and remitters (χ^2^=1.95, p=0.16)^38^. A 48-hour ADHD medication-free period was required before assessments. Parents of all participants gave informed consent following procedures approved by the London-Surrey Borders Research Ethics Committee (09/H0806/58).

**Table 1.**
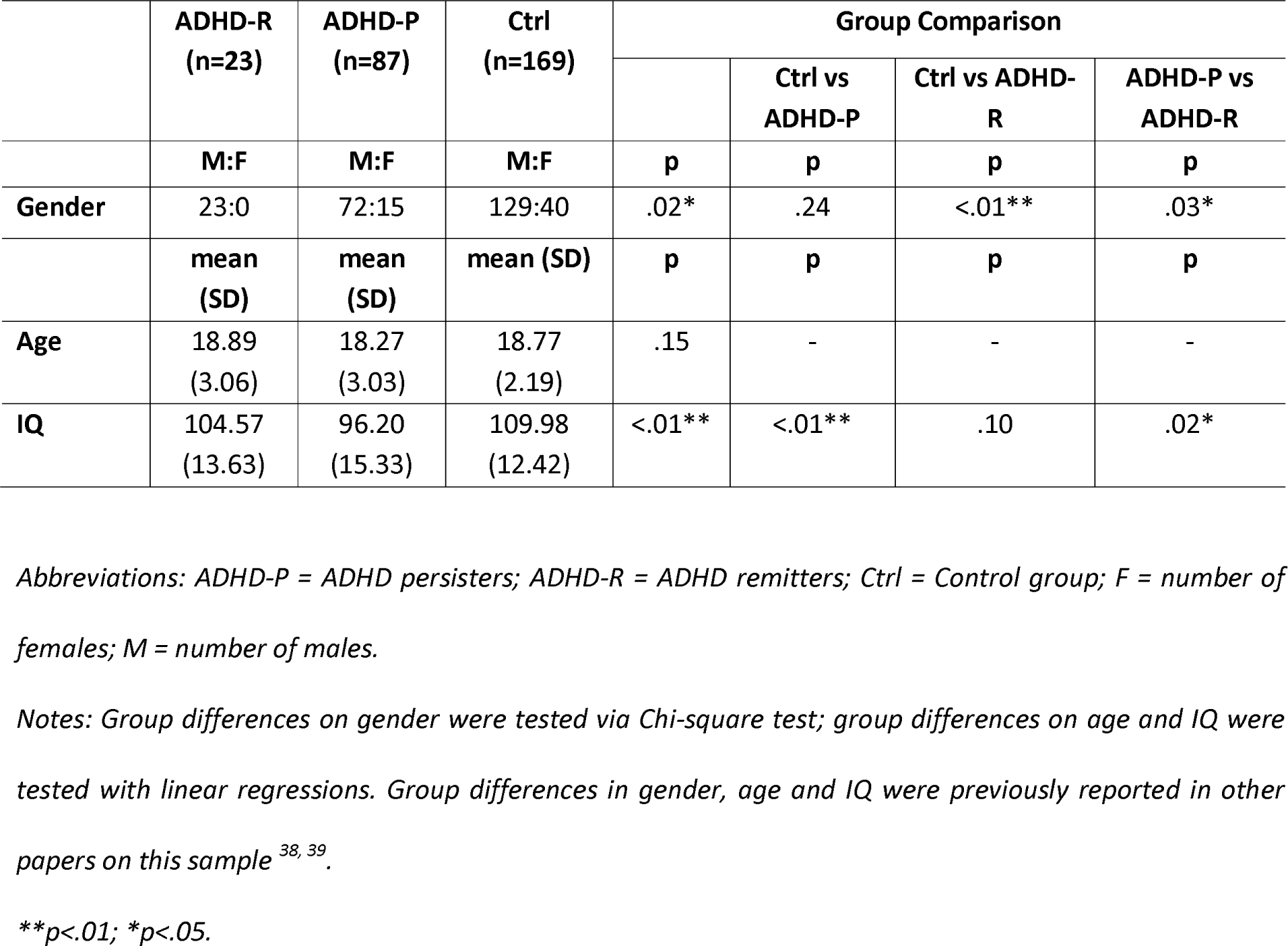
Sample demographics divided by group, with tests for differences between ADHD persisters, remitters and controls

|  | ADHD-R<br>(n=23) | ADHD-P<br>(n=87) | Ctrl<br>(n=169) | Group Comparison |  |  |  |
| --- | --- | --- | --- | --- | --- | --- | --- |
|  |  |  |  |  | Ctrl vs<br>ADHD-P | Ctrl vs ADHD-R | ADHD-P vs<br>ADHD-R |
|  | M:F | M:F | M:F | p | p | p | p |
| **Gender** | 23:0 | 72:15 | 129:40 | .02* | .24 | <.01** | .03* |
|  | mean<br>(SD) | mean<br>(SD) | mean (SD) | p | p | p | p |
| **Age** | 18.89<br>(3.06) | 18.27<br>(3.03) | 18.77<br>(2.19) | .15 | - | - | - |
| **IQ** | 104.57<br>(13.63) | 96.20<br>(15.33) | 109.98<br>(12.42) | <.01** | <.01** | .10 | .02* |

*Abbreviations: ADHD-P = ADHD persisters; ADHD-R = ADHD remitters; Ctrl = Control group; F = number of females; M = number of males.*
*Notes: Group differences on gender were tested via Chi-square test; group differences on age and IQ were tested with linear regressions. Group differences in gender, age and IQ were previously reported in other papers on this sample<sup>38, 39</sup>.*
*\*\*p<.01; \*p<.05.*

### ADHD diagnosis

The Diagnostic Interview for ADHD in Adults (DIVA)^54^ was conducted by trained researchers with parents of the ADHD probands, to assess DSM-IV-defined ADHD presence and persistence of the 18 ADHD symptoms. Evidence of impairment commonly associated with ADHD was assessed with the Barkley’s functional impairment scale (BFIS)^55^. Parent-report DIVA and impairments were used to determine ADHD status, as these were validated against objective markers (cognitive-performance and EEG measures) in this sample, whereas the same objective markers showed limited agreement with self-reported ADHD^56^. Participants were classified as “affected” at follow-up if they showed ≥6 items in either the inattention or hyperactivity-impulsivity domains on the DIVA, and ≥2 areas of impairments on the BFIS. We defined ADHD outcome using a categorical definition of persistence based on diagnoses, as well as a dimensional approach based on continuous levels of symptoms of ADHD and impairments.

### Task

The task was an adaptation of the Eriksen Flanker paradigm designed to increase cognitive load^57, 58^. In each trial a central fixation mark was replaced by a target arrow (a black 18mm equilateral triangle). Participants had to indicate whether this arrow pointed towards the left or right by pressing corresponding response buttons with their left or right index fingers. Two flanker arrows identical in shape and size to the target appeared 22mm above and below the center of the target arrow 100ms before each target. Both flankers either pointed in the same (congruent) or opposite (incongruent) direction to the target. Cognitive control and conflict monitoring are maximal during incongruent trials. When the target appeared, both target and flankers remained on the screen for 150ms, with a new trial every 1650ms. Two-hundred congruent and 200 incongruent trials were arranged in 10 blocks of 40 trials. Only incongruent trials, known to elicit greater ADHD-control differences^39, 58^, were considered in the present analysis. For further details, see Supplementary Material.

### EEG recording and processing

The EEG was recorded from a 62-channel extended 10-20 system (Brain Products, GmbH, Munich, Germany), using a 500Hz sampling-rate, impedances under 10kΩ, and recording reference at FCz. The electro-oculograms (EOGs) were recorded from electrodes above and below the left eye and at the outer canthi. Raw EEG recordings were down-sampled to 256Hz, re-referenced to the average of all electrodes (turning FCz into an active channel), and filtered using Butterworth band-pass filters (0.10-30 Hz, 24 dB/oct). All trials were visually inspected and sections containing electrical or movement artefacts were removed manually. Ocular artefacts were identified using Independent Component Analysis (ICA)^59^. Sections of data containing artefacts exceeding ±100μV or with a voltage step ≥50μV were automatically rejected. The artefact-free data were segmented in epochs between -650–1000 ms stimulus-locked to incongruent stimuli. Both trials with correct and incorrect responses were examined^39^. Only data containing ≥20 clean segments for condition were included in analyses, leaving 271 participants (83 ADHD persisters, 22 remitters, 166 controls) for correctly-responded trials and 240 (75 ADHD persisters, 20 remitters, 145 controls) for incorrectly-responded trials.

### Connectivity analysis

#### Calculation of functional connectivity

We calculated functional brain connectivity using the imaginary part of coherence (iCoh)^45, 60, 61^. This measure was chosen to ignore spurious connections between brain signals caused by volume conduction, which can substantially limit the ability to measure functional associations using EEG channels. ICoh captures the non-instantaneous connectivity between brain activities from EEG channels that are phase-lagged (i.e., delay-based)^62, 63^. Since volume conduction affects multiple scalp channels with near-zero phase delays, connectivity measured with iCoh is not contaminated by near-instantaneous artefacts of volume conduction. ICoh was measured by isolating the imaginary part of the complex number phase coherence between two signals of same frequency^45^, estimated by calculating their cross-spectrum for each time point with Fast Fourier Transforms using the EEGLAB “newcrossf” function^64^ in Matlab (The Math Works Inc., Natick, MA, USA). ICoh is measured on a scale between 0 and 1. When two signals at the same frequency have identical phase values, possibly due to volume conduction artefacts, iCoh=0. Instead, if two signals are phase lagged, iCoh>0^45^. Values of iCoh for all possible electrode pairs (62×62) were computed in the theta (4-8 Hz), alpha (8-12 Hz) and beta (12-20 Hz) bands (Figure 1), which have previously been implicated in cognitive processes engaging top-down control networks requiring coherent activity between brain areas^65-67^, such as the fronto-parietal network^68-71^.

#### Graph-theory metrics

The high multi-dimensionality of the iCoh measures was disentangled with a graph-theory approach, allowing to derive global network-based measures and describe functional associations in terms of network properties^2, 72, 73^. Graph theory is based on mathematical algorithms to quantify the relationships (“edges”) between brain signals from EEG channels, representing the “nodes” of a network. Unthresholded weighted iCoh matrices were used, in line with previous studies^6, 74-76^, where each edge is equivalent to the measured iCoh of two electrodes to preserve essential information of a network structure^2, 77, 78^. Graph-theory metrics measure the degree of network segregation (i.e., the tendency of brain regions to form local clusters with dense functional interconnections), and network integration and efficiency (i.e., the capacity of the network to become interconnected and efficiently exchange information between brain regions)^2, 79^. The following commonly-used graph measures were calculated^6, 49, 75, 77, 80^: average clustering coefficient (the probability of neighboring nodes of being inter-connected, forming densely inter-connected clusters); global efficiency (how efficient the network is in transferring information); characteristic path length and diameter (respectively, the average number of edges along the shortest paths, and the largest possible distance, between all possible pairs of nodes). Graph-theory metrics were computed separately for correctly- and incorrectly-responded trials in stimulus-locked windows, before target (pre-stimulus; -500–0 ms) and during target processing (post-stimulus; 0–500 ms) with the Brain Connectivity^47^ and BioNeCT (https://sites.google.com/site/bionectweb/home; ^3^) toolboxes.

### Statistical analyses

#### Categorical analysis based on diagnostic status

Connectivity metrics were examined with random-intercept linear models (i.e., multilevel regression models) in Stata 14 (StataCorp, College Station, TX, USA), testing for effects of group (ADHD persisters vs remitters vs controls), time window (pre-stimulus vs post-stimulus), response (correct vs incorrect) and their interaction (group-by-window-by-response). When the three-way interaction was not statistically significant, only statistically significant main effects and two-way interactions were included. For all measures, the within-group degree of change from pre-stimulus to post-stimulus was compared across groups using difference scores. All models controlled for age and took into account the degree of clustering due to family status. Cohen’s d effect sizes are presented along with test statistics, where d≥0.20 is a small effect, d≥0.50 a medium effect and d≥0.80 a large effect^81^. Given the large number of hypotheses tested, sensitivity analyses applied multiple-testing corrections with false discovery rate (FDR) on post-hoc tests with the “multproc” package, using the Simes method, which identifies those tests that remain significant^82^.

Since 80% of our sample consisted of males, but groups were not fully matched on sex (Table 1), analyses were performed on the whole sample and then repeated with females (15 ADHD persisters, 41 controls) removed. As in this sample ADHD persisters had a lower IQ than remitters^38^, and childhood IQ predicted ADHD outcome at follow-up^52^, all analyses were also re-run controlling for IQ to examine whether IQ contributes to the results. Finally, to examine brain connectivity within and between cortical regions, analyses were repeated using iCoh values within and between clusters of electrodes in different scalp regions (anterior/central/posterior) and between the two hemispheres (left/right) (for further details, see Supplementary Material).

#### Dimensional analysis with ADHD symptoms/impairment

The association between connectivity metrics and the continua of ADHD symptoms and impairment within individuals with childhood ADHD were examined with random-intercept linear models using DIVA ADHD symptom and impairment scores as independent variables, controlling for age and sex and clustering for family status. Analyses were carried out using standardized scores, thus the beta coefficients are standardized effect sizes comparable to Cohen’s d. All analyses were re-run, firstly, correcting for multiple testing, and, secondly, controlling for IQ.

#### Association between functional connectivity and cognitive performance

In an additional analysis, we examined the behavioral significance of the EEG connectivity results. We tested whether functional connectivity measured by mean iCoh was associated with task performance during the incongruent (high-conflict) condition of this task (the same task condition in which connectivity was measured). Differences between ADHD persisters, remitters and control on cognitive performance in this task can be found elsewhere^39^. Analyses were restricted to mean iCoh in the pre-stimulus window of correct trials, where differences between ADHD groups and controls were maximal based on categorical analyses. Random-intercept linear models on standardized scores tested the association of mean iCoh in theta, alpha and beta bands as independent variables with MRT, RTV and number of errors as dependent variables. These models were run separately in individuals with childhood ADHD and controls, controlling for age and sex, and clustering for family status.

## RESULTS

### Differences between ADHD persisters, remitters and controls

Post-hoc analyses revealed that, in correctly-responded trials, ADHD persisters showed greater clustering coefficient, global efficiency and mean iCoh, and lower path length and diameter compared to controls at all frequency bands in the pre-stimulus window, and only in beta in the post-stimulus windows (Table 2, Supplementary Figure 1). ADHD remitters showed lower pre-stimulus diameter in theta and beta, lower pre-stimulus path length in alpha and beta, and lower post-stimulus diameter in beta, compared to controls. ADHD remitters did not differ from persisters in any connectivity measure in correctly-responded trials, except diameter in beta (where remitters were intermediate between controls and persisters, and significantly differed from both groups) (Table 2). These findings indicate increased connectivity in both ADHD persisters and remitters compared to controls during correct trials. In error trials, group differences only emerged for clustering coefficient, global efficiency and mean iCoh in post-stimulus theta: both ADHD persisters and remitters showed reduced values in these measures (indicating lower connectivity) compared to controls, but did not differ from each other (Table 2). All three groups showed increased connectivity (greater clustering coefficient, global efficiency and mean iCoh; decreased path length and diameter) in incorrect compared to correct trials, in both pre-stimulus and post-stimulus windows (Supplementary Tables 1-2). All main and interaction effects are shown in Supplementary Table 2.

**Figure 1.**
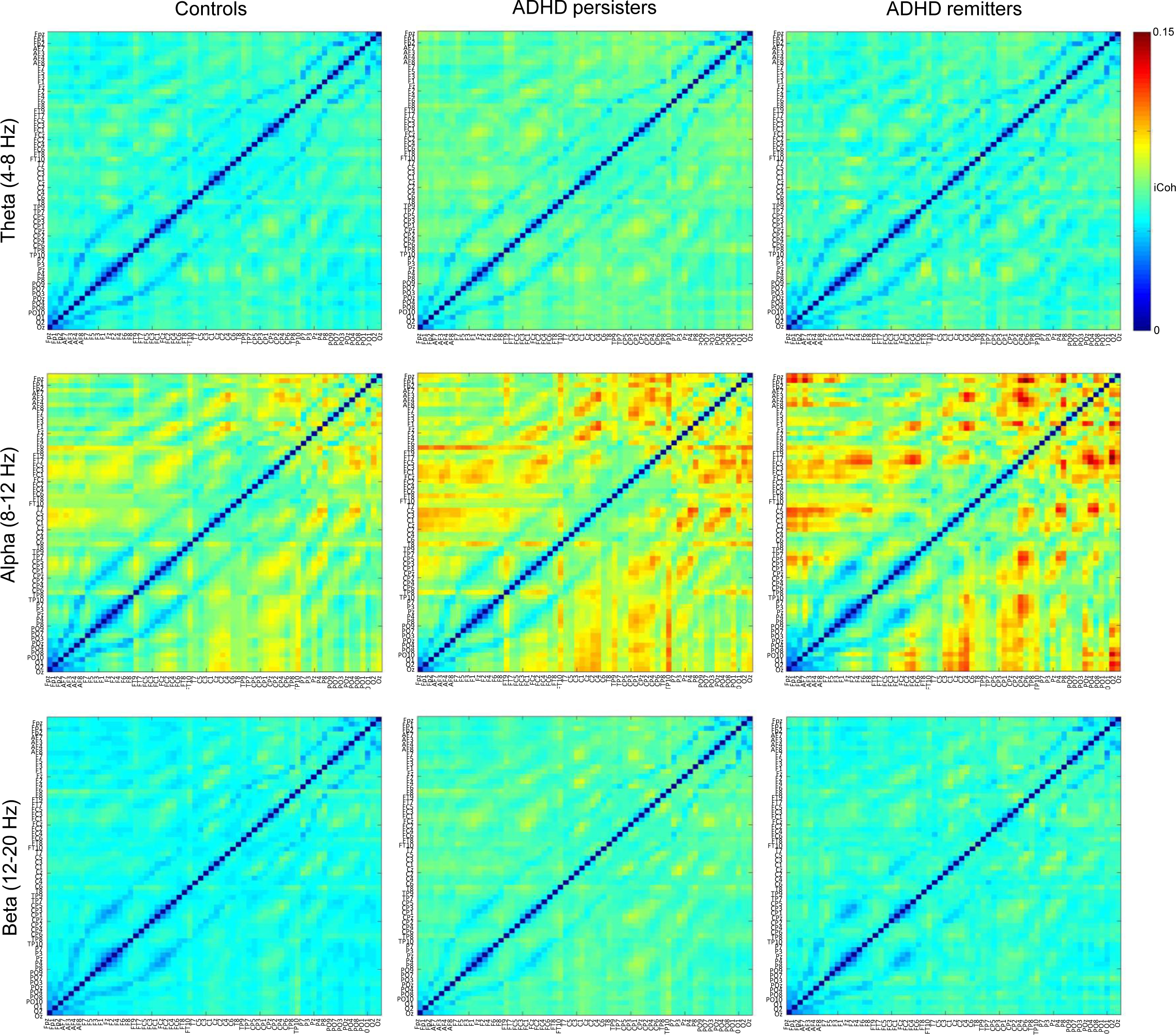
Connectivity matrices showing values of imaginary part of coherence (iCoh) in pre-stimulus theta, alpha and beta frequencies for correctly-responded trials by group (ADHD persisters, remitters and controls).

**Table 2.** Group comparisons on graph-theory and imaginary coherence measures

|  |  | Group comparison |  |  |  |  |  |  |
| --- | --- | --- | --- | --- | --- | --- | --- | --- |
| THETA |  | Overall Group | Ctrl vs ADHD-P |  | Ctrl vs ADHD-R |  | ADHD-R vs ADHD-P |  |
|  |  | p | p | d | p | d | p | d |
| Average clustering coefficient | *Pre, Corr* | 0.016* | 0.004** | 0.63 | 0.880 | 0.29 | 0.139 | 0.35 |
|  | *Pre, Err* | 0.544 | - | - | - | - | - | - |
|  | *Post, Corr* | 0.401 | - | - | - | - | - | - |
|  | *Post, Err* | <0.001*** | <0.001*** | 0.35 | 0.017* | 0.30 | 0.955 | 0.05 |
| Global efficiency | *Pre, Corr* | 0.053 | 0.019* | 0.51 | 0.901 | 0.16 | 0.145 | 0.37 |
|  | *Pre, Err* | 0.568 | - | - | - | - | - | - |
|  | *Post, Corr* | 0.189 | - | - | - | - | - | - |
|  | *Post, Err* | <0.001*** | <0.001*** | 0.35 | 0.019* | 0.30 | 0.916 | 0.05 |
| Path length | *Pre, Corr* | 0.012* | <0.001*** | 0.58 | 0.095 | 0.30 | 0.130 | 0.30 |
|  | *Pre, Err* | 0.434 | - | - | - | - | - | - |
|  | *Post, Corr* | 0.338 | - | - | - | - | - | - |
|  | *Post, Err* | 0.122 | - | - | - | - | - | - |
| Diameter | *Pre, Corr* | <0.001*** | <0.001*** | 0.64 | 0.012* | 0.49 | 0.352 | 0.17 |
|  | *Pre, Err* | 0.646 | - | - | - | - | - | - |
|  | *Post, Corr* | 0.976 | - | - | - | - | - | - |
|  | *Post, Err* | 0.279 | - | - | - | - | - | - |
| Mean imaginary coherence | *Pre, Corr* | 0.024* | 0.007** | 0.60 | 0.952 | -0.25 | 0.140 | 0.35 |
|  | *Pre, Err* | 0.562 | - | - | - | - | - | - |
|  | *Post, Corr* | 0.319 | - | - | - | - | - | - |
|  | *Post, Err* | <0.001*** | <0.001*** | 0.35 | 0.019* | 0.30 | 0.955 | 0.06 |
| ALPHA |  | Overall Group | Ctrl vs ADHD-P |  | Ctrl vs ADHD-R |  | ADHD-R vs ADHD-P |  |
|  |  | p | p | d | p | d | p | d |
| Average clustering | *Pre, Corr* | 0.001** | <0.001*** | 0.44 | 0.097 | 0.42 | 0.636 | 0.06 |
| coefficient | *Pre, Err* | 0.415 | - | - | - | - | - | - |
|  | *Post, Corr* | 0.328 | - | - | - | - | - | - |
|  | *Post, Err* | 0.084 | - | - | - | - | - | - |
| Global efficiency | *Pre, Corr* | 0.003** | 0.002** | 0.32 | 0.054 | 0.39 | 0.976 | 0.04 |
|  | *Pre, Err* | 0.325 | - | - | - | - | - | - |
|  | *Post, Corr* | 0.816 | - | - | - | - | - | - |
|  | *Post, Err* | 0.152 | - | - | - | - | - | - |
| Path length | *Pre, Corr* | <0.001*** | <0.001*** | 0.32 | 0.005** | 0.47 | 0.539 | 0.13 |
|  | *Pre, Err* | 0.709 | - | - | - | - | - | - |
|  | *Post, Corr* | 0.201 | - | - | - | - | - | - |
|  | *Post, Err* | 0.235 | - | - | - | - | - | - |
| Diameter | *Corr* | <0.001*** | <0.001*** | 0.41 | 0.054 | 0.30 | 0.610 | 0.13 |
|  | *Err* | 0.444 | - | - | - | - | - | - |
| Mean imaginary coherence | *Pre, Corr* | 0.001** | <0.001*** | 0.40 | 0.073 | 0.39 | 0.684 | 0.04 |
|  | *Pre, Err* | 0.341 | - | - | - | - | - | - |
|  | *Post, Corr* | 0.501 | - | - | - | - | - | - |
|  | *Post, Err* | 0.064 | - | - | - | - | - | - |
| BETA |  | Overall Group | Ctrl vs ADHD-P |  | Ctrl vs ADHD-R |  | ADHD-R vs ADHD-P |  |
|  |  | p | p | d | p | d | p | d |
| Average clustering coefficient | *Corr* | <0.001*** | <0.001*** | 0.79 | 0.097 | 0.51 | 0.101 | 0.31 |
|  | *Err* | 0.135 | - | - | - | - | - | - |
| Global efficiency | *Corr* | <0.001*** | <0.001*** | 0.73 | 0.137 | 0.44 | 0.098 | 0.31 |
|  | *Err* | 0.154 | - | - | - | - | - | - |
| Path length | *Corr* | <0.001*** | <0.001*** | 0.76 | 0.004** | 0.52 | 0.090 | 0.27 |
|  | *Err* | 0.343 | - | - | - | - | - | - |
| Diameter | *Corr* | <0.001*** | <0.001*** | 0.83 | 0.003** | 0.53 | 0.044* | 0.31 |
|  | *Err* | 0.221 | - | - | - | - | - | - |
| Mean imaginary coherence | *Corr* | <0.001*** | <0.001*** | *0.77* | 0.097 | *0.49* | 0.101 | 0.31 |
|  | *Err* | 0.135 | - | - | - | - | - | - |

*Abbreviations: ADHD-P = ADHD persisters; ADHD-R = ADHD remitters; Corr = trials with correct responses; Ctrl = Control group; d = Cohen's d effect size; Err = trials with incorrect responses; p = random-intercept linear model significance testing; Pre = pre-stimulus time window; Post = post-stimulus time window.*
*Notes: Random-intercept linear models tested for main effects of group (ADHD remitters vs ADHD persisters vs controls), time window (pre-stimulus vs post-stimulus) ad response (correctly- vs incorrectly-responded trials), two-way interactions (group-by-window, group-by-response, time window-by-response), and three-way interactions (group-by-window-by-response) on connectivity measures. Full results are presented in Supplementary Table 2. Since neither diameter in the alpha band, nor any measures in the beta band showed a significant group-by-window interaction, post-hoc effects of group were tested for with correctly- and incorrectly-responded trials collapsed across pre-stimulus and post-stimulus time windows. Post-hoc comparisons between groups were run only on measures showing a significant overall group effect. Age was also included as a covariate of no interest in all analyses. Data in correctly-responded trials were available for 83 ADHD persisters, 22 remitters, 166 controls; and in incorrectly-responded trials for 75 ADHD persisters, 20 remitters, 145 controls.*
*\*p<0.05; \*\*p<0.01; \*\*\*p<0.001. $d \geq 0.20$ = small effect size, $d \geq 0.50$ = medium effect (in italics) and $d \geq 0.80$ = large effect size (in bold).*

Among measures showing significant group-by-window interactions (all in theta, all except diameter in alpha, none in beta; Supplementary Table 2), significant within-group differences in changing from pre-stimulus to post-stimulus windows emerged in all groups for all theta connectivity measures, in controls only for clustering coefficient, path length and mean iCoh in the alpha band, and in both ADHD groups for global efficiency in alpha (Table 3). ADHD persisters and remitters exhibited a significantly lower degree of change than controls in all measures of theta connectivity, but no differences emerged between the two ADHD groups (Table 3).

**Table 3.** Within- and between-group effects on measures of change between pre-stimulus and post-stimulus windows in graph-theory and imaginary coherence measures

|  |  | Within-Group Change |  |  | Between-Group Change |  |  |  |  |  |
| --- | --- | --- | --- | --- | --- | --- | --- | --- | --- | --- |
|  |  | Ctrl | ADHD-P | ADHD-R | Ctrl vs ADHD-P |  | Ctrl vs ADHD-R |  | ADHD-R vs ADHD-P |  |
| THETA |  | p | p | p | p | d | p | d | p | d |
| Average clustering coefficient | *Corr* | <0.001*** | <0.001*** | <0.001*** | 0.001** | 0.42 | 0.010* | 0.41 | 0.981 | 0.05 |
|  | *Err* | <0.001*** | <0.001*** | <0.001*** | 0.011* | 0.33 | 0.618 | 0.06 | 0.370 | 0.26 |
| Global efficiency | *Corr* | <0.001*** | <0.001*** | <0.001*** | 0.002** | 0.40 | 0.014* | 0.38 | 0.997 | 0.04 |
|  | *Err* | <0.001*** | <0.001*** | <0.001*** | 0.017* | 0.31 | 0.643 | 0.06 | 0.400 | 0.25 |
| Path length | *Corr* | <0.001*** | <0.001*** | <0.001*** | <0.001*** | 0.61 | 0.014* | 0.44 | 0.506 | 0.19 |
|  | *Err* | <0.001*** | <0.001*** | <0.001*** | 0.058 | 0.27 | 0.776 | 0.11 | 0.209 | 0.36 |
| Diameter | *Corr* | <0.001*** | <0.001*** | <0.001*** | <0.001*** | 0.61 | 0.016* | 0.43 | 0.499 | 0.19 |
|  | *Err* | <0.001*** | <0.001*** | <0.001*** | 0.058 | 0.20 | 0.776 | 0.14 | 0.209 | 0.33 |
| Mean imaginary coherence | *Corr* | <0.001*** | <0.001*** | <0.001*** | 0.001** | 0.42 | 0.011* | 0.40 | 0.995 | 0.05 |
|  | *Err* | <0.001*** | <0.001*** | <0.001*** | 0.013* | 0.33 | 0.632 | 0.06 | 0.378 | 0.26 |
| ALPHA |  | Ctrl | ADHD-P | ADHD-R | Ctrl vs ADHD-P |  | Ctrl vs ADHD-R |  | ADHD-R vs ADHD-P |  |
|  |  | p | p | p | p | d | p | d | p | d |
| Average clustering coefficient | *Corr* | 0.002** | 0.910 | 0.767 | 0.055 | 0.27 | 0.091 | 0.38 | 0.704 | 0.09 |
|  | *Err* | 0.001** | 0.981 | 0.599 | 0.069 | 0.28 | 0.267 | 0.16 | 0.468 | 0.13 |
| Global efficiency | *Corr* | 0.728 | 0.004** | 0.045* | 0.071 | 0.27 | 0.147 | 0.40 | 0.705 | 0.11 |
|  | *Err* | 0.155 | 0.029* | 0.683 | 0.019* | 0.38 | 0.140 | 0.25 | 0.389 | 0.15 |
| Path length | *Corr* | 0.002** | 0.856 | 0.319 | 0.124 | 0.20 | 0.049* | 0.42 | 0.349 | 0.23 |
|  | *Err* | 0.011* | 0.831 | 0.931 | 0.023* | 0.37 | 0.094 | 0.33 | 0.743 | 0.07 |
| **Mean imaginary coherence** | *Corr* | 0.020* | 0.491 | 0.472 | 0.064 | 0.27 | 0.111 | 0.37 | 0.735 | 0.08 |
|  | *Err* | 0.001** | 0.545 | 0.791 | 0.015* | 0.40 | 0.087 | 0.30 | 0.469 | 0.13 |

*Abbreviations: ADHD-P = ADHD persisters; ADHD-R = ADHD remitters; Corr = trials with correct responses; Ctrl = Control group; d = Cohen's d effect size; Err = trials with incorrect responses; p = random-intercept linear model significance testing.*
*Notes: Random-intercept linear models tested for main effects of group (ADHD remitters vs ADHD persisters vs controls), time window (pre-stimulus vs post-stimulus) ad response (correctly- vs incorrectly-responded trials), two-way interactions (group-by-window, group-by-response, time window-by-response), and three-way interactions (group-by-window-by-response) on connectivity measures. Full results are presented in Supplementary Table 2. Post-hoc tests on within- and between-group effects of change were run only on measures showing a significant group-by-window interaction. Since this interaction was not significant in diameter in the alpha band or in any measures in the beta band, post-hoc within- and between-groups effects of change were not tested. Age was also included as a covariate of no interest in all analyses. Data in correctly-responded trials were available for 83 ADHD persisters, 22 remitters, 166 controls; and in incorrectly-responded trials for 75 ADHD persisters, 20 remitters, 145 controls. $d \geq 0.20$ = small effect size, $d \geq 0.50$ = medium effect (in italics), $d \geq 0.80$ = large effect.*
*\* $p < 0.05$ ; \*\* $p < 0.01$ ; \*\*\* $p < 0.001$ .*

Multiple-testing corrections (controlling the FDR at 15%) on post-hoc group comparisons (separately for ADHD persisters vs controls, ADHD remitters vs controls, ADHD persisters vs remitters) showed that all statistically significant differences between controls and ADHD remitters, and between controls and ADHD persisters remained significant. The only previously significant difference between ADHD persisters and remitters (in beta diameter) was no longer significant when correcting for multiple testing. All significant group differences on measures of pre-stimulus/post-stimulus change remained significant after correcting for multiple testing.

All results remained unchanged when rerunning analyses on the male-only sample (Supplementary Tables 3-4), except that the p-values of certain tests that were statistically significant in the full sample became trend-level effects (p=0.05-0.10). All effect sizes were similar to those on the full sample, suggesting that these non-significant results may be due to lower power in this smaller sample.

Results of group comparisons on connectivity measures in pre- and post-stimulus were largely unchanged when IQ was included as a covariate in categorical analyses (Supplementary Table 5). A few differences between persisters and controls on measures of pre-stimulus/post-stimulus change in theta and alpha connectivity during error trials were no longer significant (Supplementary Table 6).

Results of analyses on group differences in local connectivity within cortical regions (within anterior/central/posterior regions and within left/right hemispheres) and these three cortical regions and two hemispheres, were consistent with those on whole-brain connectivity (for full results, see Supplementary Material).

### Association with ADHD symptoms and impairment

In dimensional analyses on participants with childhood ADHD, no association emerged between ADHD symptoms and any connectivity measure in theta, alpha or beta frequencies in correct or error trials (Table 4). Functional impairment was not associated with any connectivity measure in the theta band, but showed associations with a subgroup of measures in alpha and beta in correct and error trials (Table 4). Results remained largely unchanged when controlling for IQ (Supplementary Table 7). Statistically significant associations that emerged with ADHD impairment were no longer significant after applying multiple-testing corrections.

**Table 4.** Dimensional associations between graph-theory and imaginary coherence measures and interview-based DIVA ADHD symptom counts and clinical impairment within the ADHD group only
(n=110), controlling for age and gender

| THETA |  | ADHD symptoms |  | Impairment |  |
| --- | --- | --- | --- | --- | --- |
| | | $\beta$ (95% CIs) | p | $\beta$ (95% CIs) | p |
| **Average clustering coefficient** | *Pre, Corr* | -0.005 (-0.202;0.193) | 0.964 | 0.160 (-0.065;0.384) | 0.163 |
|  | *Pre, Err* | 0.020 (-0.177;0.216) | 0.844 | 0.178 (-0.040;0.398) | 0.110 |
|  | *Post, Corr* | -0.021 (-0.218;0.174) | 0.827 | 0.176 (-0.0041;0.393) | 0.111 |
|  | *Post, Err* | -0.088 (-0.259;0.084) | 0.315 | -0.068 (-0.263;0.127) | 0.494 |
| **Global efficiency** | *Pre, Corr* | -0.037 (-0.216;0.142) | 0.685 | 0.089 (-0.115;0.292) | 0.393 |
|  | *Pre, Err* | 0.004 (-0.181;0.189) | 0.969 | 0.167 (-0.280;0.373) | 0.110 |
|  | *Post, Corr* | -0.043 (-0.267;0.152) | 0.667 | 0.145 (-0.074;0.365) | 0.194 |
|  | *Post, Err* | -0.111 (-0.279;0.057) | 0.196 | -0.152 (-0.344;0.040) | 0.120 |
| **Path length** | *Pre, Corr* | 0.033 (-0.145;0.211) | 0.716 | -0.123 (-0.322;0.076) | 0.226 |
|  | *Pre, Err* | -0.027 (-0.231;0.178) | 0.797 | -0.175 (-0.402;0.053) | 0.132 |
|  | *Post, Corr* | 0.067 (-0.140;0.273) | 0.528 | -0.100 (-0.332;0.131) | 0.395 |
|  | *Post, Err* | 0.108 (-0.092;0.308) | 0.290 | 0.056 (-0.171;0.282) | 0.630 |
| **Diameter** | *Pre, Corr* | 0.049 (-0.135;0.232) | 0.601 | -0.109 (-0.321;0.102) | 0.310 |
|  | *Pre, Err* | -0.043 (-0.251;0.165) | 0.685 | -0.168 (-0.107;0.028) | 0.153 |
|  | *Post, Corr* | 0.030 (-0.161;0.222) | 0.756 | -0.116 (-0.329;0.098) | 0.287 |
|  | *Post, Err* | 0.100 (-0.100;0.300) | 0.328 | 0.020 (-0.202;0.242) | 0.861 |
| **Mean imaginary coherence** | *Pre, Corr* | 0.013 (-0.205;0.180) | 0.898 | 0.142 (-0.077;0.361) | 0.204 |
|  | *Pre, Err* | 0.015 (-0.178;0.208) | 0.878 | 0.175 (-0.040;0.390) | 0.110 |
|  | *Post, Corr* | -0.028 (-0.224;0.168) | 0.778 | 0.167 (-0.051;0.385) | 0.134 |
|  | *Post, Err* | -0.096 (-0.266;0.074) | 0.268 | -0.101 (-0.295;0.093) | 0.306 |
| ALPHA |  | ADHD symptoms |  | Impairment |  |
| | | $\beta$ (95% CIs) | p | $\beta$ (95% CIs) | p |
| **Average clustering coefficient** | *Pre, Corr* | 0.014 (-0.186;0.215) | 0.894 | 0.044 (-0.186;0.274) | 0.708 |
|  | *Pre, Err* | 0.051 (-0.130;0.233) | 0.578 | 0.157 (-0.049;0.363) | 0.135 |
|  | *Post, Corr* | 0.064 (-0.121;0.249) | 0.500 | 0.256 (0.056;0.456) | 0.012* |
|  | *Post, Err* | 0.117 (-0.063;0.297) | 0.204 | 0.223 (0.000;0.001) | 0.034* |
| **Global efficiency** | *Pre, Corr* | -0.027 (-0.231;0.176) | 0.794 | 0.091 (-0.326;0.145) | 0.450 |
|  | *Pre, Err* | 0.033 (-0.160;0.226) | 0.738 | 0.130 (-0.089;0.349) | 0.245 |
|  | *Post, Corr* | 0.052 (-0.125;0.230) | 0.563 | 0.199 (0.004;0.394) | 0.046* |
|  | *Post, Err* | 0.108 (-0.063;0.280) | 0.216 | 0.222 (0.021;0.419) | 0.031* |
| **Path length** | *Pre, Corr* | <0.001 (-0.184;0.183) | 0.998 | 0.080 (-0.129;0.289) | 0.452 |
|  | *Pre, Err* | -0.026 (-0.219;0.167) | 0.793 | -0.132 (-0.347;0.083) | 0.229 |
|  | *Post, Corr* | -0.052 (-0.237;0.132) | 0.580 | -0.202 (-0.404;0.000) | 0.050 |
|  | *Post, Err* | -0.131 (-0.319;0.057) | 0.172 | -0.222 (-0.436;-0.008) | 0.042* |
| **Diameter** | *Pre, Corr* | -0.027 (-0.223;0.129) | 0.784 | -0.051 (-0.277;0.175) | 0.659 |
|  | *Pre, Err* | -0.080 (-0.272;0.113) | 0.417 | -0.177 (-0.397;0.043) | 0.114 |
|  | *Post, Corr* | -0.053 (-0.245;0.139) | 0.588 | -0.232 (-0.442;-0.023) | 0.030* |
|  | *Post, Err* | -0.134 (-0.332;0.064) | 0.185 | -0.201 (-0.428;0.027) | 0.083 |
| **Mean imaginary coherence** | *Pre, Corr* | 0.003 (-0.202;0.195) | 0.973 | -0.003 (-0.229;0.224) | 0.981 |
|  | *Pre, Err* | 0.049 (-0.152;0.251) | 0.631 | 0.165 (-0.063;0.393) | 0.156 |
|  | *Post, Corr* | 0.062 (-0.121;0.245) | 0.505 | 0.242 (0.045;0.441) | 0.016* |
|  | *Post, Err* | 0.115 (-0.063;0.294) | 0.204 | 0.223 (0.018;0.427) | 0.033* |
| **BETA** |  | **ADHD symptoms** |  | **Impairment** |  |
|  |  | **β (95% CIs)** | **p** | **β (95% CIs)** | **p** |
| **Average clustering coefficient** | *Pre, Corr* | 0.124 (-0.100;0.349) | 0.278 | 0.306 (-0.062;0.550) | 0.014* |
|  | *Pre, Err* | 0.051 (-0.149;0.252) | 0.613 | 0.203 (-0.022;0.429) | 0.077 |
|  | *Post, Corr* | 0.093 (-0.141;0.328) | 0.435 | 0.264 (0.001;0.527) | 0.049* |
|  | *Post, Err* | 0.045 (-0.160;0.250) | 0.666 | 0.166 (-0.062;0.393) | 0.153 |
| **Global efficiency** | *Pre, Corr* | 0.100 (-0.119;0.319) | 0.372 | 0.299 (-0.061;0.536) | 0.014* |
|  | *Pre, Err* | 0.047 (-0.155;0.248) | 0.650 | 0.210 (-0.016;0.436) | 0.069 |
|  | *Post, Corr* | 0.068 (-0.167;0.302) | 0.572 | 0.248 (-0.015;0.512) | 0.065 |
|  | *Post, Err* | 0.042 (-0.163;0.248) | 0.688 | 0.166 (-0.062;0.394) | 0.152 |
| **Path length** | *Pre, Corr* | -0.092 (-0.289;0.105) | 0.361 | -0.233 (-0.450;-0.016) | 0.035* |
|  | *Pre, Err* | -0.067 (-0.264;0.131) | 0.508 | -0.198 (-0.419;0.023) | 0.080 |
|  | *Post, Corr* | -0.073 (-0.280;0.134) | 0.490 | -0.191 (-0.425;0.043) | 0.110 |
|  | *Post, Err* | -0.080 (-0.283;0.124) | 0.444 | -0.163 (-0.387;0.061) | 0.153 |
| **Diameter** | *Pre, Corr* | -0.118 (-0.318;0.083) | 0.251 | -0.253 (-0.474;0.033) | 0.024* |
|  | *Pre, Err* | -0.094 (-0.301;0.112) | 0.372 | -0.166 (-0.395;0.063) | 0.157 |
|  | *Post, Corr* | -0.105 (-0.312;0.102) | 0.320 | -0.190 (-0.425;0.045) | 0.114 |
|  | *Post, Err* | -0.089 (-0.299;0.122) | 0.410 | -0.124 (-0.357;0.108) | 0.294 |
| **Mean imaginary coherence** | *Pre, Corr* | 0.118 (-0.105;0.341) | 0.301 | 0.305 (-0.063;0.548) | 0.013* |
|  | *Pre, Err* | 0.051 (-0.150;0.251) | 0.620 | 0.207 (-0.019;0.432) | 0.072 |
|  | *Post, Corr* | 0.085 (-0.150;0.321) | 0.478 | 0.259 (-0.005;0.523) | 0.054 |
|  | *Post, Err* | 0.044 (-0.162;0.249) | 0.676 | 0.166 (-0.062;0.394) | 0.153 |

Abbreviations: $\beta$ = standardized regression coefficient; CIs = confidence intervals; Corr = trials with correct responses; Err = trials with incorrect responses; p = random-intercept linear model significance testing; Pre = pre-stimulus time window; Post = post-stimulus time window. Data in correctly-responded trials were available for 105 childhood ADHD participants (83 ADHD persisters, 22 remitters); and in incorrectly-responded trials for 95 childhood ADHD participants (75 ADHD persisters, 20 remitters).
Notes: Random-intercept linear models tested for the effect of ADHD symptom count/impairment on each connectivity measure. $\beta \geq 0.20$ = small effect size, $\beta \geq 0.50$ = medium effect, $\beta \geq 0.80$ = large effect.
\* $p < 0.05$ .

### Association with cognitive performance

Greater mean iCoh in the theta and beta bands showed statistically significant effects on greater RTV in the childhood ADHD group, and on greater number of errors in both childhood ADHD and control groups (Supplementary Table 8). Alpha mean iCoh was not significantly associated with any performance measure in either group. None of the pre-stimulus iCoh connectivity measures had a significant effect on MRT in either group.

## DISCUSSION

Using a network-based EEG functional connectivity approach, our results indicate that ADHD persisters show widespread over-connectivity underlying cognitive-control processes compared to controls, as well as reduced adjustments of connectivity with changed task demands. ADHD remitters showed similar impairments as persisters, and differed from controls in most measures of connectivity and connectivity adjustments. These findings indicate that hyper-connectivity and reduced ability to modulate connectivity patterns with task demands characterize adolescents and young adults with both persistent and remitted ADHD. Atypical functional connectivity during cognitive-control processes may thus represent an enduring deficit in adolescents and adults with childhood ADHD, irrespective of their diagnostic status at follow up.

Two main connectivity impairments emerged in individuals with persistent ADHD compared to controls. Firstly, ADHD persisters showed increased global connectivity (higher iCoh), network segregation (higher clustering coefficient), efficiency (higher global efficiency) and integration (lower path length and diameter) at all frequency bands prior to target onset in trials with correct behavioral responses, as well as during target processing in beta oscillations. The increases in functional connectivity are consistent with a previous EEG study reporting pre-target over-connectivity in children with ADHD^30^, and more generally supports evidence indicating hyper-connectivity in ADHD during task performance^20, 23, 28, 29^. Connectivity in theta, alpha and beta oscillations during cognitive tasks is associated with cognitive processes engaging control networks and requiring coordination of activity between distributed brain areas^65-67^. Here, over-connectivity in these frequency ranges in persistent ADHD may reflect exaggerated interactions between brain regions, both during the inactive pre-stimulus period and during cognitive target processing. Considering the high cognitive demands induced by incongruent stimuli in this highly effortful task, which requires a response at every trial, increased connectivity may reflect hyper-connectivity in distributed brain networks underlying higher-level cognitive functions. Secondly, while all groups showed increased theta connectivity in changing from pre-stimulus to post-stimulus windows, this change was reduced in ADHD persisters compared to controls. This result in individuals with ADHD may point to a reduced ability to modulate brain connectivity patterns in slow oscillations from a relatively inactive context to a condition requiring cognitive engagement. This finding is in line with previous reports indicating reduced regulation of brain activity in ADHD between different cognitive states^83-85^. Overall, these findings show widespread connectivity impairments underlying cognitive-control processes in ADHD persisters, and advance our understanding of the neural underpinnings of persistent ADHD in adolescence and early adulthood.

Our study represents the first investigation into EEG connectivity in adolescents and adults with remitted ADHD. In several connectivity measures sensitive to impairments in persisters, ADHD remitters were impaired compared to controls and indistinguishable from persisters, consistent with our hypotheses. ADHD remitters also showed the same reduction in all measures of pre-stimulus/post-stimulus change in theta connectivity displayed by persisters. As such, brain connectivity impairments were insensitive to ADHD outcome (remission/persistence) in adolescence and early adulthood, and may represent enduring deficits irrespective of current diagnostic status. Findings from dimensional analyses supported these results, as most connectivity measures were unrelated to continuous levels of ADHD symptoms and impairments in participants with childhood ADHD. Of note, while results of categorical analyses were largely unchanged after correcting for multiple testing, the few significant associations between connectivity and functional impairment (all with small effect sizes) did not survive multiple-testing corrections. Overall, these connectivity findings in remitters are consistent with previous cognitive-EEG studies, including our previous analyses on this sample^38, 39^, reporting that executive-functioning measures were insensitive to ADHD outcomes in adolescence and adulthood^35-39^. They also partially align with results from a previous resting-state connectivity fMRI study, which found over-connectivity in remitters compared to controls and no differences between remitters and persisters^42^. A clinical implication is that connectivity impairments underlying executive-control processes may not be suitable targets for interventions for ADHD, consistent with previous evidence of no effects of stimulants on EEG connectivity in ADHD^25, 86^. Future EEG studies should examine whether connectivity during less effortful activities, such as rest or non-executive processes, represent markers of remission, similar to cognitive-EEG measures of non-executive processes in our previous studies^38-40^.

Of note, while widespread group differences emerged in correctly-responded trials, group differences in error trials, likely representing a failure of cognitive control, emerged only in three measures of post-stimulus theta connectivity. The limited group differences in incorrect responses may suggest that a suboptimal pattern of brain connectivity may attenuate the differences in brain-network profiles between neurotypical individuals and individuals with ADHD, who are prone to making more incorrect responses in this task^39^. In addition, all groups showed greater connectivity before and during incorrect responses than correct responses. In an additional analysis testing whether hyper-connectivity was related to impairments in cognitive performance during this task, we found that pre-stimulus hyper-connectivity in theta and beta oscillations was associated with fewer correct responses in individuals with childhood ADHD and controls, and with increased RTV in individuals with childhood ADHD. A suboptimal pattern of hyper-connectivity underlying cognitive control processes may thus lead to dysfunctional behavioral responses, both in neurotypical individuals and in individuals with childhood ADHD.

A limitation of this study is that, despite the large sample, the low ADHD remission rate at follow-up resulted in a relatively small group of remitters. Therefore, we could not exclude the possibility that some non-significant group differences could be due to low power. However, the moderate effect sizes (d=0.38-0.53) between ADHD remitters and controls, but negligible or small (d=0.02-0.36) between remitters and persisters, in measures showing ADHD persister-control differences suggest that we had sufficient power to detect, with the current sample sizes, differences in connectivity with at least moderate effect sizes. In addition, our sample included young adults as well as adolescents, who are still undergoing rapid cortical maturation. While analyses controlled for age, future follow-up assessments with participants having reached adulthood could provide further insight into developmental patterns. Finally, while the current EEG connectivity analyses allowed precise temporal resolution and connectivity estimates unaffected by volume-conduction artefacts, the relatively poor spatial resolution of scalp-EEG did not allow precise localization of the brain networks. Yet, results of local connectivity within and between cortical regions were consistent with those of whole-brain analyses, indicating comparable effects in more localized networks.

In conclusion, we report new evidence of shared atypical connectivity profiles in adolescents and young adults with persistent and remitted ADHD. These connectivity alterations may represent enduring deficits and neural signatures associated with having a history of childhood ADHD, but unrelated to current diagnostic status. Connectivity impairments underlying executive processes may represent associated characteristics or risk factors in ADHD^87^, which do not follow the developmental pathways of clinical profiles. Future studies should explore the presence of potential compensatory mechanisms in individuals with remitted ADHD that enable developmental improvements in clinical profiles and non-executive cognitive processes^38-40^, despite persistence of enduring connectivity alterations.

## AKNOWLEDGMENTS

We thank all who made this research possible: our participants, their families and research workers Jessica Deadman, Hannah Collyer and Sarah-Jane Gregori. This project was supported by generous grants from Action Medical Research and the Peter Sowerby Charitable Foundation (GN1777) to Prof Jonna Kuntsi. Initial cognitive assessments of the ADHD and control groups in childhood, and the recruitment of the control sample were supported by UK Medical Research Council grant (G0300189) to Prof Jonna Kuntsi. Initial sample recruitment of the ADHD group was supported by NIMH grant (R01MH062873) to Prof Stephen V. Faraone. Dr Giorgia Michelini was supported by a 1+3 PhD studentship awarded by the MRC Social, Genetic and Developmental Psychiatry Centre, Institute of Psychiatry, Psychology and Neuroscience, King’s College London (G9817803), and by a Short-term fellowship by the European Molecular Biology Organization (EMBO ASTF 218-2015). Prof Philip Asherson is supported by generous grants from the National Institute for Health Research (NIHR) Biomedical Research Centre for Mental Health at King’s College London, Institute of Psychiatry, Psychology and Neuroscience and South London and Maudsley National Health Service (NHS) Foundation Trust. Dr Ioannis Bakolis is supported by the NIHR Biomedical Research Centre at South London and Maudsley NHS Foundation Trust and by the NIHR Collaboration for Leadership in Applied Health Research and Care South London at King’s College Hospital NHS Foundation Trust. This paper represents independent research part-funded by the NIHR Biomedical Research Centre at South London and Maudsley NHS Foundation Trust and King’s College London. The views expressed are those of the authors and not necessarily those of the NHS, the NIHR or the Department of Health and Social Care.

